# Epistatic effects between amino acid insertions and substitutions mediate toxin-resistance of vertebrate Na^+^,K^+^-ATPases

**DOI:** 10.1101/2022.09.01.506169

**Authors:** Shabnam Mohammadi, Halil İbrahim Özdemir, Pemra Ozbek, Fidan Sumbul, Josefin Stiller, Yuan Deng, Andrew J. Crawford, Hannah M. Rowland, Jay F. Storz, Peter Andolfatto, Susanne Dobler

## Abstract

The recurrent evolution of resistance to cardiotonic steroids (CTS) across diverse animals most frequently involves convergent amino-acid substitutions to the H1-H2 extracellular loop of Na^+^,K^+^-ATPase (NKA). Previous work established that hystricognath rodents (e.g. chinchilla) and pterocliform birds (sandgrouse) have convergently evolved amino-acid insertions in the H1-H2 loop, but their functional significance is not known. Using protein engineering, we show that these insertions have distinct effects on CTS resistance of NKA in the two lineages that strongly depend on intramolecular interactions with other residues. Removing the insertion in the chinchilla lineage unexpectedly increases CTS resistance and decreases NKA activity. In the sandgrouse lineage, the insertion works in concert with the substitution Q111R to increase CTS resistance while maintaining wild-type ATPase activity levels. Molecular docking simulations provide additional insight into the biophysical mechanisms responsible for the context-specific CTS insensitivity of the enzyme. Our results highlight the diversity of genetic substrates that underlie CTS insensitivity in vertebrate NKA and reveal how amino-acid insertions can alter the phenotypic effects of point mutations at key sites in the same protein domain.

## Introduction

The evolution of toxin resistance in animals is among the best studied examples of adaptive molecular evolution (Brodie 2009). In many cases, diverse animals have convergently evolved resistance to the same toxin, allowing one to examine the extent to which genetic background and other factors constrain the process of adaptive protein evolution (McGlothlin et al. 2016; Mohammadi et al. 2022). Recent studies have highlighted how the potentially adaptive effects of particular amino acid mutations can strongly depend on the protein sequence background on which they arise (Weinreich et al. 2006; Gong and Bloom 2014; Storz 2018). Substantially less attention has been paid to the importance of insertions and deletions on these intramolecular constraints. While insertions and deletions are relatively rare, they nonetheless harbor substantial potential to impact the properties and evolvability of proteins (de la Chaux et al. 2007). Here, we investigate the impact of convergently evolved insertions in the context of the evolution of cardiotonic steroids (CTS) resistance in animals.

CTS comprise a diverse group of plant- and animal-derived secondary compounds that are often used as a means of chemical defense against herbivores and predators (Krenn and Kopp 1998; Hutchinson et al. 2007; Agrawal et al. 2012). CTS are toxic to animals because they inhibit Na^+^,K^+^-ATPase (NKA), a heterodimeric transmembrane protein that consists of a catalytic α-subunit, encoded by members of the ATP1A gene family, and a glycoprotein β-subunit, encoded by members of the ATP1B gene family (Fig. 1A; (Köksoy 2002; Aperia 2007). NKAs play a critical role in maintaining membrane potential via trans-membrane exchange of Na^+^ and K^+^ ions and are consequently vital for the maintenance of diverse physiological functions including neural signal transduction, muscle contraction, and cellular homeostasis (Blanco and Mercer 1998; Mobasheri et al. 2000; Bagrov et al. 2009). CTSs bind to a specific domain of the α-subunit of NKA (ATP1A), and binding affinity is strongly determined by the H1-H2 extracellular loop, which forms part of that domain (Fig. 1A; (Laursen et al. 2015).

**Figure 1.**
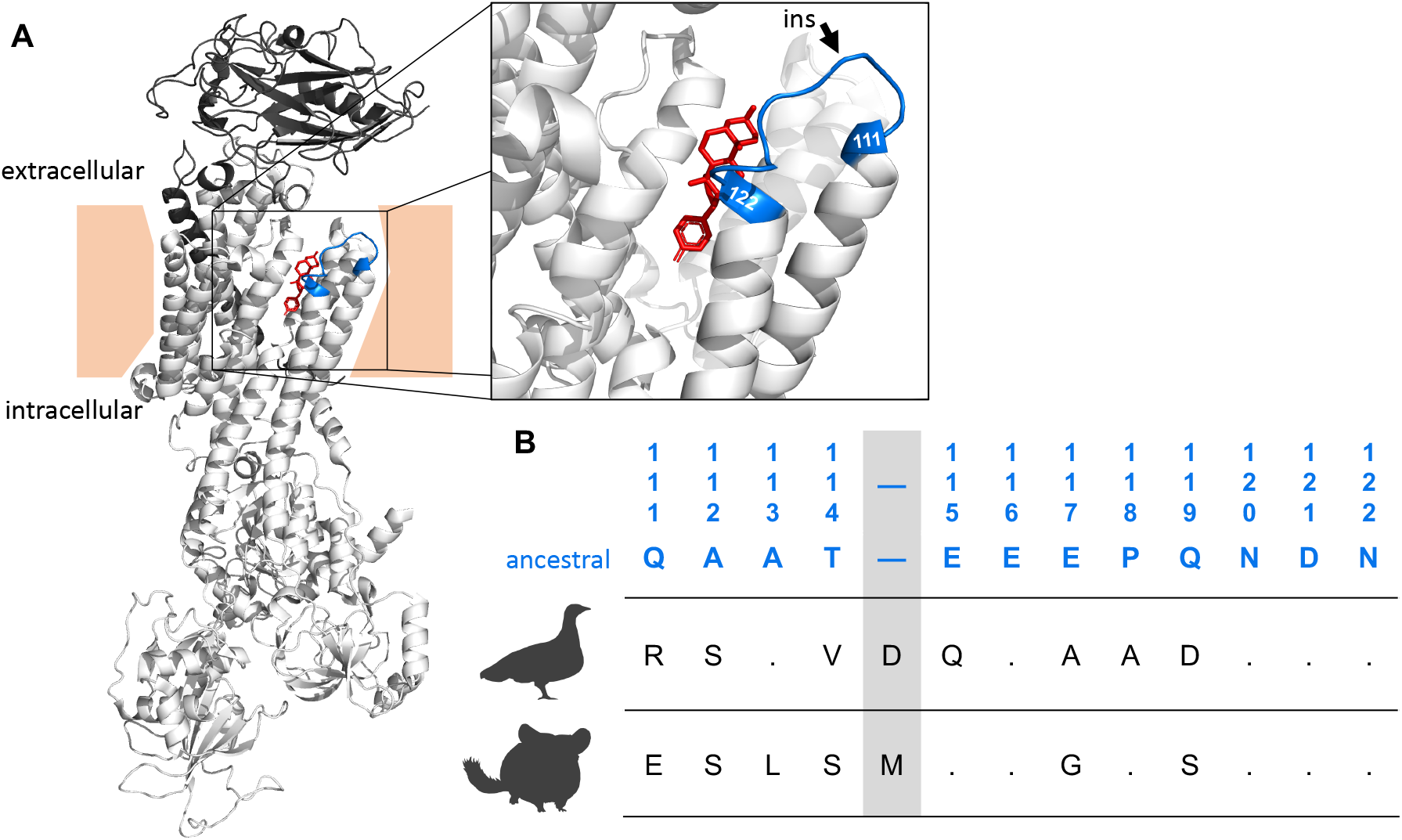
(A) Crystal structure of an Na^+^,K^+^-ATPase with a bound cardiotonic steroid (bufalin) in red (*Sus scrofa*; PDB 4RES). The α-subunit is colored in light grey tones and the β-subunit is colored in dark grey. The zoomed-in panel shows the H1-H2 extracellular loop, highlighted in blue. Two sites, 111 and 122, at which substitutions have been repeatedly implicated in CTS resistance are labeled in blue. (B) The ancestral tetrapod amino acid sequence of the H1-H2 extracellular loop (Mohammadi et al. 2022) is indicated by blue text and the numbering follows the sheep ATP1A1 sequence convention. The corresponding wildtype sequences for the yellow-throated sandgrouse (*Pterocles gutturalis)* and the common chinchilla (*Chinchilla lanigera*) are listed below. The inferred position of the insertions in both species are highlighted in grey.

Convergent evolution of resistance to CTS in diverse animals is often mediated, at least in part, by the evolution of NKA ‘target-site insensitivity’. Experimental studies have revealed a surprising consistency in the underlying molecular mechanisms of CTS insensitivity of NKA (Ujvari et al. 2015; Storz 2016; Karageorgi et al. 2019; Taverner et al. 2019; Mohammadi et al. 2021; Mohammadi et al. 2022). Different combinations of amino-acid substitutions have been reported in animals that have evolved CTS-resistance, and available evidence suggests that some substitutions impede CTS binding by preventing the formation of hydrogen bonds between hydroxyl groups of CTS and polar residues of the H1-H2 extracellular loop (Laursen et al. 2015). In tetrapods, different isoforms of the NKA α-subunit are encoded by three paralogous members of a multigene family (ATP1A1-ATP1A3), with a fourth paralog specific to mammals (ATP1A4). Resistance-conferring amino acid substitutions have identified in paralogs A1-A3, but are most common in ATP1A1, which is also the most ubiquitously expressed (Ujvari et al. 2013; Ujvari et al. 2015; Mohammadi et al. 2016; Marshall et al. 2018; Mohammadi et al. 2022).

In taxa as diverse as milkweed-feeding insects and toad-eating vertebrates, the evolution of CTS resistance is often associated with amino acid substitutions at sites 111 and 122 in the H1-H2 extracellular loop of ATP1A (Price and Lingrel 1988; Dobler et al. 2012; Zhen et al. 2012; Groen and Whiteman 2021; Mohammadi et al. 2021; Mohammadi et al. 2022). Consequently, these two sites have been the focus of experimental efforts to characterize molecular mechanisms of target-site insensitivity in NKA. However, results from several recent studies have revealed nonadditive interactions (i.e., intramolecular epistasis) involving sites 111, 122, and other sites in ATP1A that impinge on multiple aspects of protein function and affect resistance to CTS (Karageorgi et al. 2019; Taverner et al. 2019; Mohammadi et al. 2021; Mohammadi et al. 2022). Two recent studies reported a two-amino acid insertion in addition to multiple amino acid substitutions in the H1-H2 loop of pyrgomorphid grasshopper species that feed on CTS-defended plants (Dobler et al. 2019; Yang et al. 2019). Together, the insertion and substitutions at sites 111 and 122 were shown to result in a strong increase in CTS resistance when added to the wild-type *Drosophila melanogaster* ATP1A ortholog (Dobler et al. 2019).

Similar insertions were recently documented in two vertebrate lineages, mammals (hystricognath rodents) and birds (sandgrouse) (Mohammadi et al. 2022). While initial site-directed mutagenesis experiments on the ATP1A1 of these two taxa focused only on substitutions at sites 111 and 122 of ATP1A1, they yielded puzzling results (Mohammadi et al. 2022). Specifically, the wildtype ATP1A1 of chinchilla (*Chinchilla lanigera,* a representative of hystricognath rodents) does not exhibit CTS resistance. However, engineering the replacement N122D (which introduces an amino-acid state known to confer CTS resistance in ATP1A1 of other tetrapod species) (Price et al. 1990; Mohammadi et al. 2021; Mohammadi et al. 2022) confers orders of magnitude higher CTS resistance than that observed in other rodents with the same substitution. Conversely, the wildtype ATP1A1 of the sandgrouse, (*Pterocles gutturalis,* Aves: Pterocliformes) exhibits substantial resistance. However, mutational reversion of the derived “resistant” R111 to the ancestral “sensitive” amino-acid state (Q111) did not diminish resistance of the sandgrouse ATP1A1, suggesting that determinants of resistance lie elsewhere in the protein (Mohammadi et al. 2022). We hypothesize that the proximate insertions observed in hystricognath rodents and sandgrouse potentially impact resistance to CTS, but may do so in different ways in these two lineages.

We have recently established that the effects of resistance-conferring substitutions at 111 and 122 strongly depend on the protein sequence background in which they occur and that these context-dependent effects likely depend on a small number of sites (Mohammadi et al. 2022). The dependence on a limited number of sites may account for the observation of convergent CTS resistance substitutions observed among highly divergent taxa. In support of this view, Mohammadi et al. (Mohammadi et al. 2021) showed that negative pleiotropic effects caused by substitutions at 111 and 122 can be rescued by 10 (or fewer) of the 19 amino acid differences distinguishing the backgrounds of CTS-resistant and sensitive ATP1A1 paralogs of grass frogs. We therefore asked whether the observed insertions in chinchilla and sandgrouse ATP1A1 are important contributors to background-dependent effects. To answer this question, we used site-directed mutagenesis to test whether these insertions alter the effect of point mutations at sites 111 and 122 known to confer NKA target-site insensitivity.

## Results

### Origin of the H1-H2 loop insertions

A previous survey of ATP1A1 of 117 mammals and 70 birds established that chinchilla (a hystricognath rodent) and sandgrouse (a pterocliform bird) have convergently evolved amino-acid insertions between positions 114 and 115 of the H1-H2 extracellular loop (Mohammadi et al. 2022). In the chinchilla, the inserted amino acid is methionine (insM), and in the sandgrouse it is aspartic acid (insD). To infer the evolutionary origins of the insertions, we separately estimated maximum likelihood phylogenies of 26 rodent ATP1A1 sequences and 22 avian ATP1A1 sequences obtained from publicly available sources (Table S1; Supplementary Datasets 1-2). The estimated phylogeny of rodent ATP1A1 sequences indicates that the insertion in the H1-H2 loop evolved in a recent common ancestor of the hystricognath rodents, which includes chinchillas, porcupines, guinea pigs, nutrias, mole rats, and allies (Marivaux et al. 2004) (Fig. 2A). This implies that the insM is ancient, dating from 36 to 39 million years ago (Sallam et al. 2009). The estimated phylogeny of avian ATP1A1 sequences indicates that the insertion evolved in the common ancestor of Pterocliformes, which includes all sandgrouse species (Fig. 2B). Despite its relatively restricted phylogenetic distribution, insM of sandgrouse may also be quite ancient given that the common ancestor of Pterocliformes diverged from its sister group, Mesitornithiformes, between 45 and 55 million years ago (Kuhl et al. 2021). In both cases the insertions are flanked by multiple additional amino acid substitutions.

**Figure 2.**
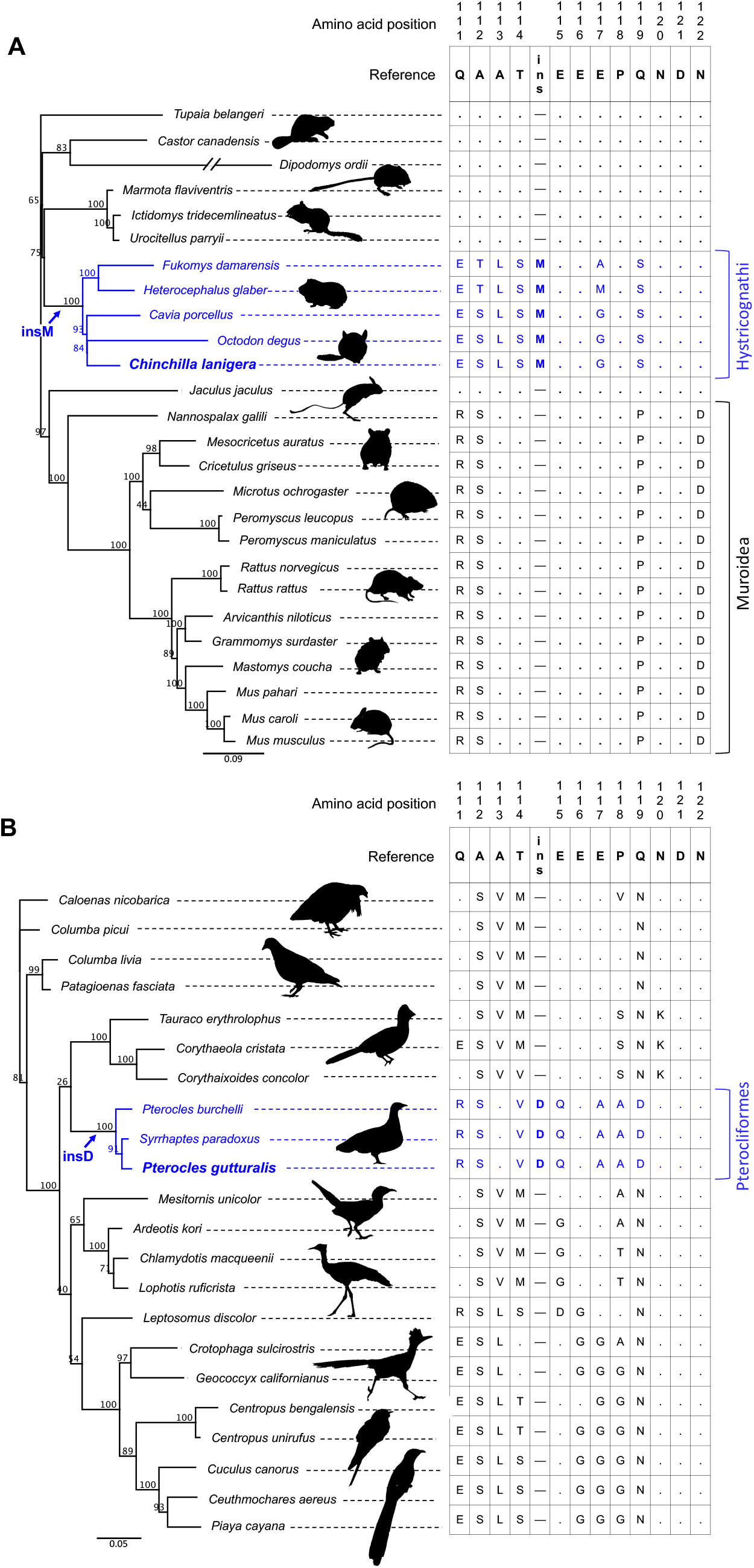
Maximum likelihood phylogenies inferred from ATP1A1 nucleotide sequences using IQ-TREE v 2.1.2 (Minh et al. 2020). Amino acid sequences of the H1-H2 loop (positions 111 to 122) are aligned to the right of each phylogeny. (A) Rodent protein tree inferred from an alignment of 26 protein-coding DNA sequences. Branch-tip labels in blue denote species with the insM insertion and includes all members of the clade Hystricognathi. All five species also share several other amino acid substitutions, indicating that they originated in the common ancestor of Hystricognathi. (B) Bird tree inferred from an alignment of 22 protein-coding DNA sequences. Branch-tip labels in blue denote species with the insD insertion, which include all sampled sandgrouse (Pterocliformes). Similar to the hystricognath rodents, the insertion (insD in this case) is accompanied by several other substitutions shared by all sandgrouse.

### The effect of insertions on CTS resistance

To test the functional effects of the H1-H2 loop insertions, we used protein engineering to evaluate the effects of various combinations of amino acid substitutions and insertions (Table 1). In the case of chinchilla, we deleted insM on the wildtype ATP1A1 and introduced a known resistance-conferring mutation at site 122 (N122D) on backgrounds that did or did not include insM. We chose D122 because this amino-acid state is shared by all members of the closely related clade of murid rodents (Price and Lingrel 1988; Mohammadi et al. 2022); Fig. 2A). In the case of sandgrouse, we deleted insD on the wildtype ATP1A1 and reverted a known resistance-conferring substitution at site 111 (R111Q) (Price and Lingrel 1988; Mohammadi et al. 2021) on backgrounds that did or did not include insD.

**Table 1.**
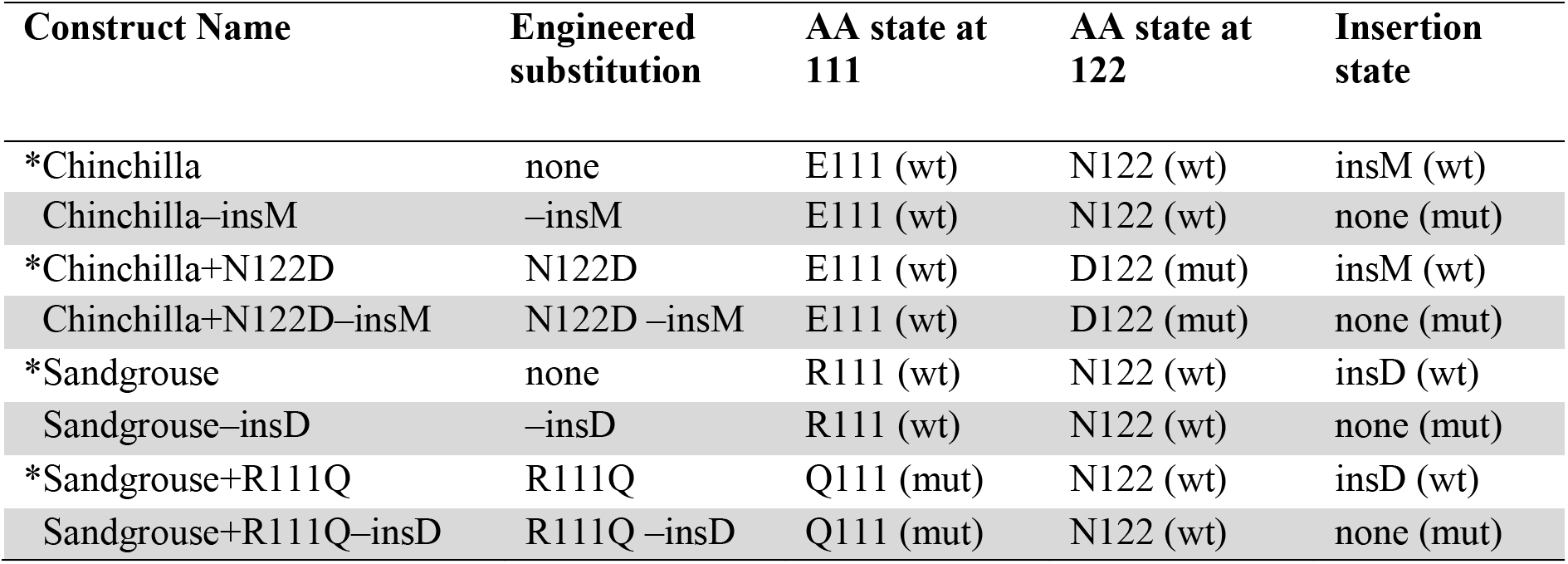
List of gene constructs used to test functional effects of amino acid mutations and species-specific insertions in ATP1A1 of the common chinchilla (*Chinchilla lanigera*) and yellow-throated sandgrouse (*Pterocles gutturalis*). The corresponding wildtype ATP1B1 gene of each recombinant protein construct was co-expressed with ATP1A1. Asterisks indicate constructs generated by (Mohammadi et al. 2022). The abbreviations “wt” and “mut” indicate wildtype or derived states, respectively.

| Construct Name | Engineered substitution | AA state at 111 | AA state at 122 | Insertion state |
| --- | --- | --- | --- | --- |
| *Chinchilla | none | E111 (wt) | N122 (wt) | insM (wt) |
| Chinchilla–insM | –insM | E111 (wt) | N122 (wt) | none (mut) |
| *Chinchilla+N122D | N122D | E111 (wt) | D122 (mut) | insM (wt) |
| Chinchilla+N122D–insM | N122D –insM | E111 (wt) | D122 (mut) | none (mut) |
| *Sandgrouse | none | R111 (wt) | N122 (wt) | insD (wt) |
| Sandgrouse–insD | –insD | R111 (wt) | N122 (wt) | none (mut) |
| *Sandgrouse+R111Q | R111Q | Q111 (mut) | N122 (wt) | insD (wt) |
| Sandgrouse+R111Q–insD | R111Q –insD | Q111 (mut) | N122 (wt) | none (mut) |

For each recombinant protein, we quantified the level of CTS resistance as IC_50_, which is the molar concentration of CTS needed to reduce protein activity by 50%. The rate of ATP hydrolysis in the absence of CTS was used as a measure of native protein function (protein activity; Table S2; Fig. 3). For ATP1A1 of both chinchilla and sandgrouse, we measured IC_50_ of the wildtype proteins and a combinatorially complete set of single- and double-mutant genotypes (Fig. 3; Table S3). Surprisingly, removing insM from the wildtype chinchilla ATP1A1 caused a significant increase in IC_50_ (Tukey HSD, p = 0.007), suggesting that the insertion confers NKA with greater CTS sensitivity. As expected, adding N122D onto the wildtype chinchilla ATP1A1 caused a significant increase in IC_50_ (Tukey HSD, p < 0.001), however, adding N122D in the absence of insM had no effect on IC_50_. A significant interaction between insM and N122D (2-way ANOVA interaction term, F_1,8_=145.4, p = 2e-6; Table S3) further indicates that these directionally contrasting effects are due to epistatic interactions involving the insertion and site 122 within the chinchilla ATP1A1.

**Figure 3.**
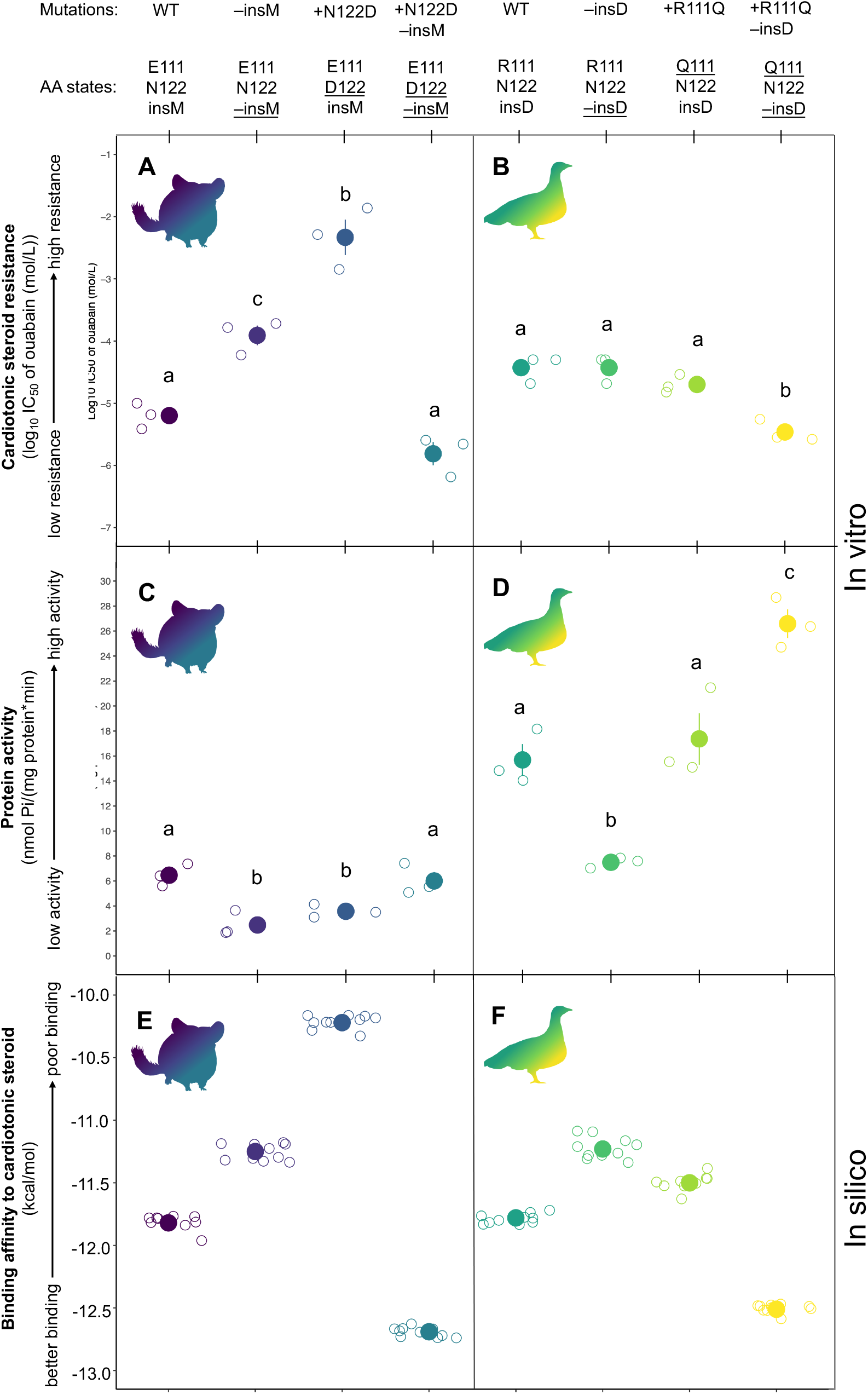
Joint in vitro functional properties of eight engineered Na^+^,K^+^-ATPases (NKAs) from the common chinchilla (*Chinchilla lanigera*; panels A and C) and yellow-throated sandgrouse (*Pterocles gutturalis*; panels B and D). For each recombinant protein, amino-acid states at positions 111, 122, and the insertion are denoted, and mutagenesis-derived changes at these states are underlined. For each species, recombinant proteins consist of (from left to right), the wildtype NKA, the wildtype NKA–insert, the wildtype NKA+mutation, and the wildtype NKA+mutation–insert. Mean ± SEM log_10_ IC_50_ (i.e., CTS resistance) of three biological replicates for each recombinant NKA is plotted on the y-axis of panels A and B. Mean ± SEM ATP hydrolysis rate (i.e., protein activity) for the same proteins is plotted on the y-axis of panels C and D. Raw data from three biological replicates of each protein are shown as open circles and jittered with respect to the x-axis. Significant differences (pairwise *t* test) between proteins for each panel are indicated by different letters. Corresponding in silico docking scores from the best docking position of ouabain docked in the common chinchilla (panel E) and yellow-throated sandgrouse (panel F) recombinant NKAs. The best docking member for each case is defined as the docking position closest to the position of ouabain in the published co-crystal structure (PDB id: 7DDJ). The predicted binding energy of 10 individual docking simulations is represented by open circles, jittered with respect to the x-axis, and the mean values (± SEM) by colored spheres.

In the case of sandgrouse, removing insD from wildtype ATP1A1 has no effect on IC_50_. Further, introducing the reversion of a well-documented resistance substitution, R111Q also results in no change in IC_50_. However, introducing R111Q in the absence of insD does result in a significant decrease in IC_50_ (Tukey HSD, p < 0.001). These results indicate that both insD and Q111R have the potential to confer resistance, but that the two substitutions jointly exhibit “diminishing-returns” epistasis (2-way ANOVA interaction term, F_1,8_=11.6, p = 0.009; Table S3).

### The effect of insertions on NKA activity

Experiments on the various chinchilla and sandgrouse NKA mutants also reveal strong epistatic effects on protein activity. The direction of these effects changed depending on the mutation combination. In chinchilla, removing insM from the wildtype chinchilla ATP1A1 significantly reduced protein activity (Tukey HSD, p = 0.004). Adding N122D to wildtype ATP1A1 also reduced protein activity (Tukey HSD, p = 0.024) but adding N122D to ATP1A1 in the absence of insM did not alter activity. In line with the nonadditive effects we observed, we found a significant interaction between insM and N122D on protein activity (2-way ANOVA interaction term, F_1,8_=34.57, p <0.001; Table S3). Overall, protein activity in chinchilla ATP1A1 decreases as resistance increases (Fig. 3C-D).

Experiments on the sandgrouse ATP1A1 demonstrate similar multi-directional epistatic effects. Removing insD from the wildtype sandgrouse ATP1A1 resulted in significantly reduced protein activity (Tukey HSD, p = 0.011). Adding the R111Q reversion substitution onto the wildtype sandgrouse ATP1A1 did not affect protein activity, however, adding R111Q in the absence of insD resulted in a significant increase in protein activity (Tukey HSD, p = 0.002). The significant interaction between insD and R111Q (2-way ANOVA interaction term, F_1,8_=42.01, p <0.001; Table S3) indicates that this pair of substitutions interact epistatically with respect to protein activity.

In both species, removing the insertions results in reduced protein activity, indicating that both are important in maintaining the enzyme’s function. To the extent that “+R111Q-insD” represents the ancestral state of the sandgrouse lineage, we infer that insD likely mitigated the large decrease in protein activity associated with the Q111R substitution. The role of insM in the chinchilla lineage may have similarly mitigated negative pleiotropic effects caused by substitutions at other derived sites on the protein.

### The biophysical mechanism of insertion effects on resistance

To investigate the biophysical mechanisms underlying the effects on resistance observed in recombinant proteins, we used a homology model of a high-affinity structure of NKA to perform molecular docking simulations (Zhen et al. 2012; Kanai et al. 2021) using Autodock Vina 1.1.2 (Trott and Olson 2010). We modeled each recombinant protein and performed docking simulations using ouabain, the CTS used in our functional experiments (see Figs. 3E-F). The trend in the docking scores for amino acid substitutions and insertions is consistent with their observed effects on CTS resistance in functional experiments. The amino acid substitutions and insertions altered the interaction network between ATP1A1 and ouabain, thereby affecting ligand-binding affinity.

In the case of chinchilla ATP1A1, removing insM from the wildype chinchilla ATP1A1 results in loss of H-bonds between the liganded ouabain and S119 and E908, despite the formation of new H-bonds with E111 and D121 (Figs. 3E; 4B; 6A; S3A). The N122D mutation in the presence of insM is predicted to alter the H-bonding network of the liganded ouabain and the receptor (Fig. 4C). This is largely attributable to the loss of an H-bond between ouabain and N122. Although new H-bonds form between C103, Y901 and ouabain, the loss of the H-bond with residue 122 is predicted to reduce binding affinity (i.e., increasing resistance; Figs. 3E, 4C, 6A). N122D in the absence of insM results in loss of H-bonds with S119 and E908 while recovering the H-bond with D122, as well as formation of additional H-bonds with E116, E111, and D121 (Figs. 3E; 4D). The only difference between the two cases lies in the H-bond between ouabain the the sidechain of E116, which seems crucial for stronger binding. Overall, across all individual modelling replicates, N122D in the absence of insM results in the highest number of H-bonds, with multiple bonds formed by residues E111, E116 and D122 (Fig. S3A), which is predicted to increase ouabain binding-affinity. The distances among H-bond donor-acceptor pairs are also shorter for the N122D mutation in the absence of insM, leading to strong to moderate interactions, which increases binding-affinity (Fig. 6A).

**Figure 4.**
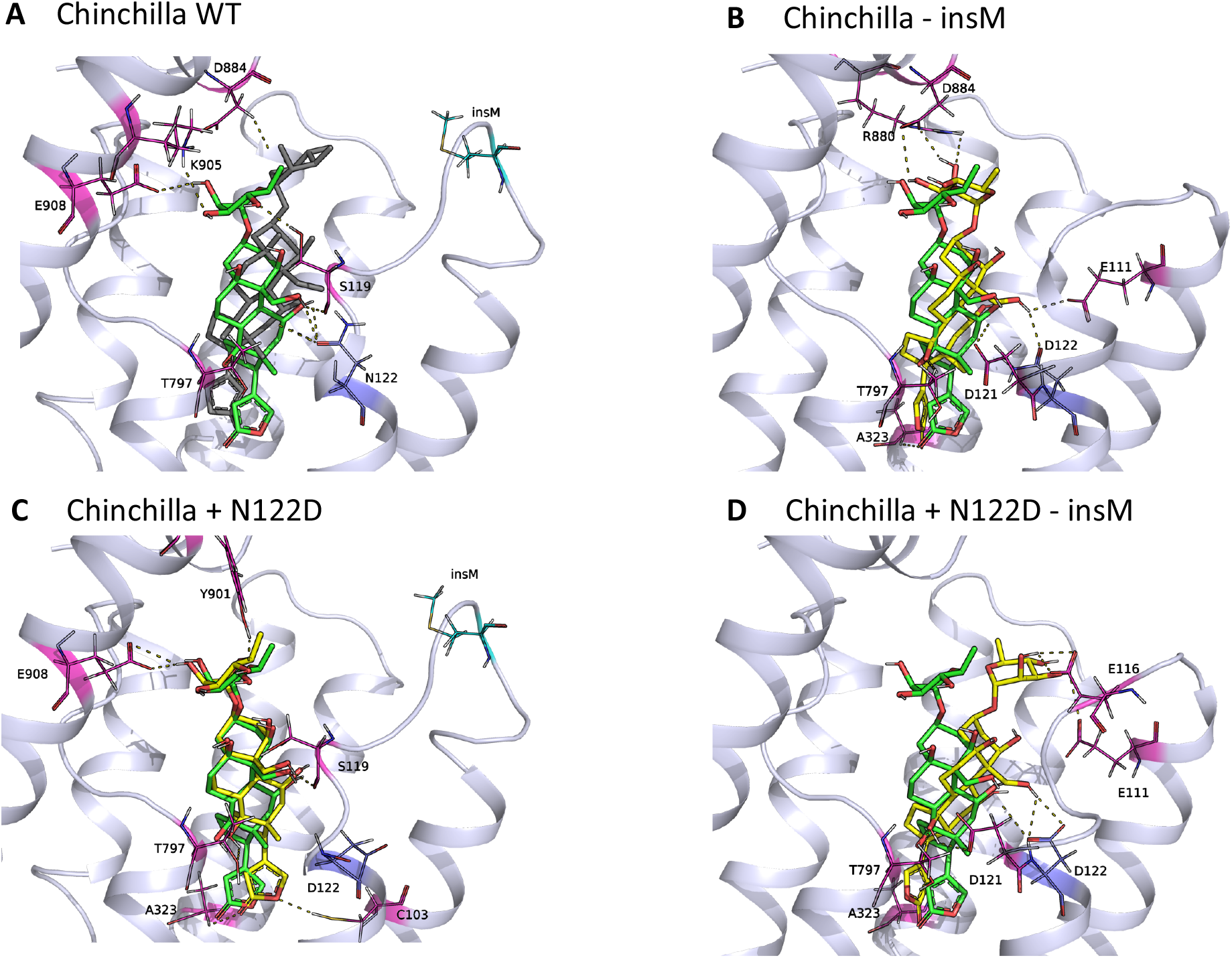
The docked structure of ouabain in the binding pocket of chinchilla ATP1A1 modeled together with hydrogen bonds obtained from molecular docking simulations. The dark gray ligand in panel A belongs to the co-crystal structure of ouabain with ATP1A1 (PDB ID: 7DDJ), and the green ligand in all panels represents docked ouabain position to the chinchilla wildtype (WT) model. The yellow ligand in panels B-D shows docked ouabain conformations in different mutant ATP1A1 structures. The interacting residues are labeled and shown in stick form and the H-bonds between the ATP1A1 and ouabain are represented as dashed lines. The transmembrane helices of the homology models are superimposed to the same region of the co-crystal structure. The extracellular region of the α-subunit is removed for simplicity.

In the case of sandgrouse ATP1A1, removal of insD from the wildtype ATP1A1 results in stronger H-bonds with D884 but weakened H-bonds at R111 and D121 without changing overall affinity (Figs. 3F; 5B; 6B). R111Q is predicted to produce a slight alteration of the H-bond network, thereby altering the docked conformation of ouabain. The R111Q mutation in the presence of insD results in the loss of an H-bond between ouabain and R111, while forming a new H-bond with K905 (Figs. 3F; 5C; S3B), but the net result is not expected to significantly alter ligand affinity. On the other hand, the R111Q substitution in the absence of insD recovers H-bonding at position 111. This leads to new H-bonds between ouabain and residues Q115 and G796 and stronger binding to ouabain (Figs. 3F; 5D; 6B; S3B). Thus, Q115 and G796 contribute significantly to the binding of oubain (Fig. 5D).

**Figure 5.**
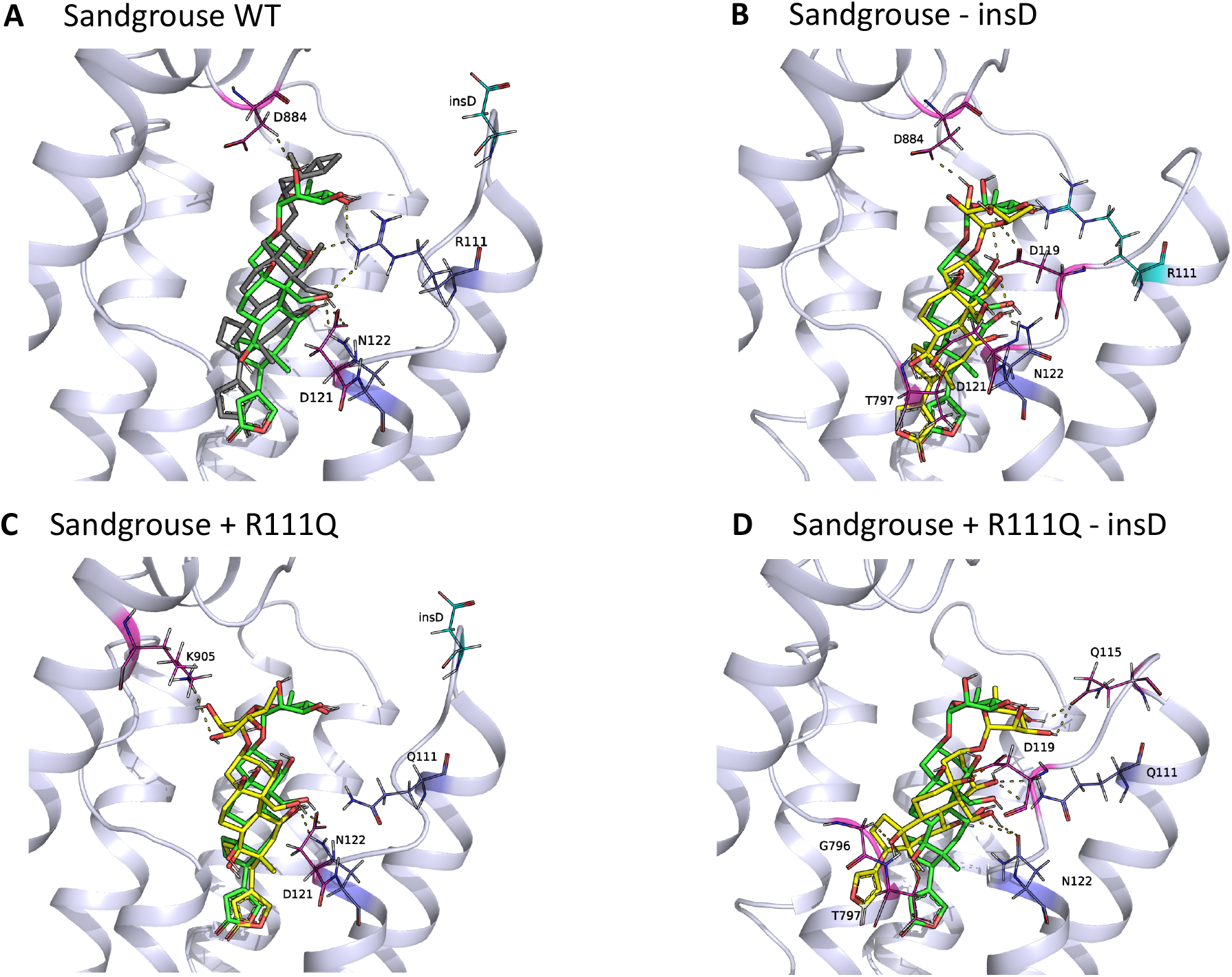
The docked structure of ouabain in the binding pocket of sandgrouse ATP1A1 modeled together with hydrogen bonds obtained from molecular docking simulations. The dark gray ligand in panel A belongs to the co-crystal structure of ouabain with ATP1A1 (PDB ID: 7DDJ), and the green ligand in all panels represents docked ouabain pose to the Sandgrouse wildtype (WT) model. The yellow ligand in panels B-D shows docked ouabain conformations in different mutant ATP1A1 structures. The interacting residues are labeled and shown in sticks and the H-bonds between the ATP1A1 and ouabain are represented as dashed lines. The transmembrane helices of the homology models are superimposed to the same region of the co-crystal structure. The extracellular region of the α-subunit is removed for simplicity.

**Figure 6.**
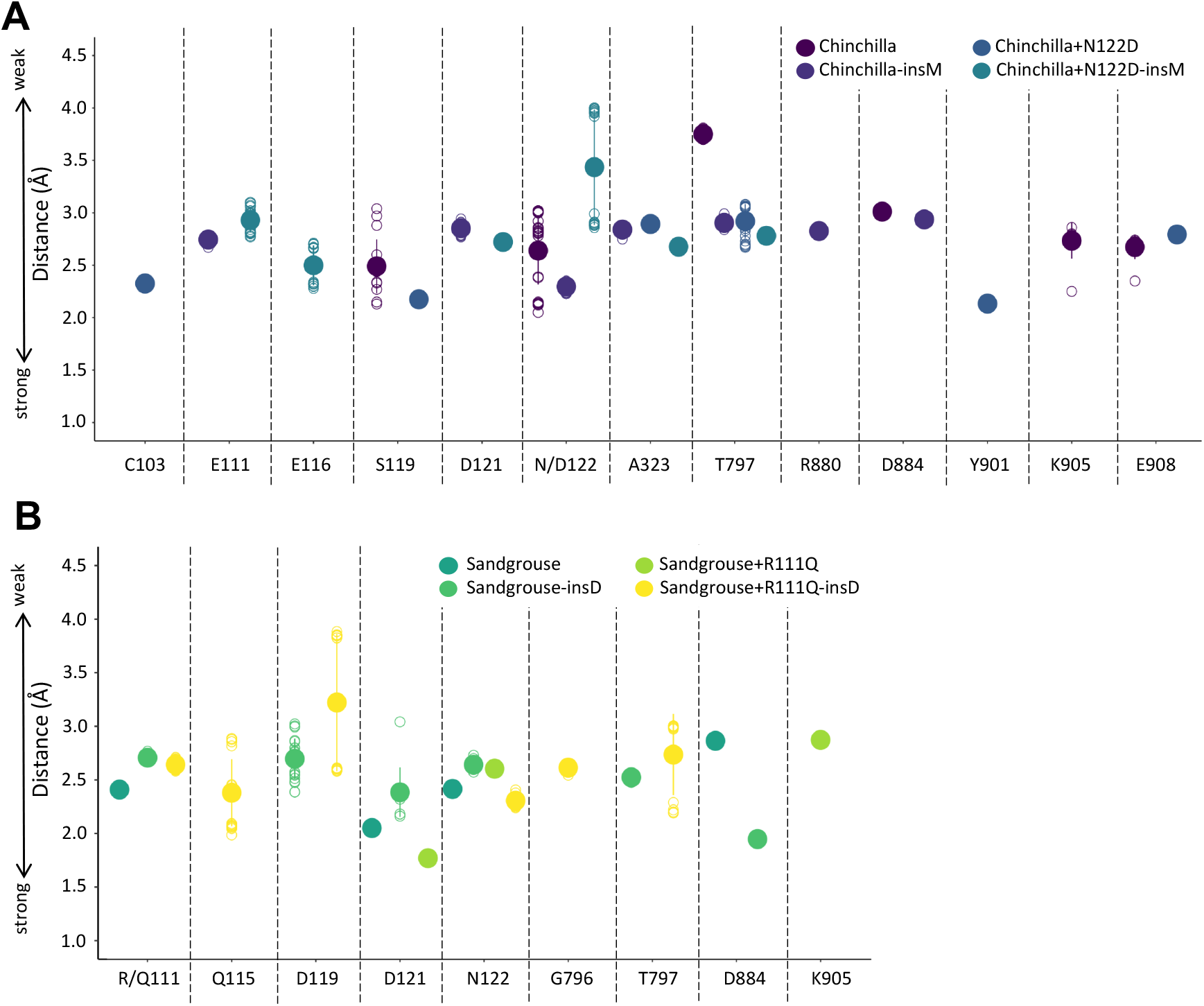
The distribution of distances between the donor-acceptor pairs making H-bonds among corresponding ATP1A1 residues and ouabain for A) chinchilla and B) sandgrouse. The distances among bond forming atoms depict the strength of corresponding H-bonds. The lower the bond distance, the stronger the bond is.

The predicted effects of amino acid substitutions and insertions follow a similar pattern for the ATP1A1 of both species. In ATP1A1 of both chinchilla and sandgrouse, when insertions are removed from the H1-H2 loop with derived amino acid states at 111 and 122, respectively, the binding affinity is strengthened, resulting in less CTS-resistant enzymes. Conversely, when the insertions are removed from the wildtype ATP1A1 of both species, the binding affinity is weakened resulting in increased CTS resistance.

## Discussion

Previous work suggests that the evolution of CTS resistance has been constrained by epistatic effects of resistance-conferring substitutions and that these constraints are likely mediated by a few key sites on the protein (Karageorgi et al. 2019; Taverner et al. 2019; Mohammadi et al. 2021; Mohammadi et al. 2022). Our protein engineering experiments and molecular docking simulations of chinchilla and sandgrouse NKA corroborate this pattern and reveal functional effects and epistatic interactions involving insertions in the H1-H2 extracellular loop of ATP1A1. Previous work also suggests that resistance-conferring substitutions of the H1-H2 loop are often associated with negative pleiotropy on overall protein function (Zhen et al. 2012; Dalla et al. 2017; Taverner et al. 2019; Yang et al. 2019; Mohammadi et al. 2021; Mohammadi et al. 2022) and that these effects are often mitigated by other amino acid substitutions throughout the protein (Karageorgi et al. 2019; Taverner et al. 2019; Mohammadi et al. 2021). Our mutagenesis experiments reveal that, while amino acid substitutions at sites 111 and 122 can compromise NKA activity, proximate amino acid insertions can play a role in mitigating these effects. In the following paragraphs, we discuss how the results of our mutagenesis experiments and models contribute to our understanding of the evolution of target-site insensitivity of NKA to CTS and the potential importance of insertion-deletion substitutions to protein function and evolution.

N122D is a derived substitution shared by all Muroid rodents (Muroidea) and neotropical grass frogs (*Leptodactylus),* and is always associated with the substitution Q111R (Mohammadi et al. 2022). While N122D has been shown to confer resistance individually in ATP1A1 of mouse (Price and Lingrel 1988) and neotropical grass frog (Mohammadi et al. 2021), the protein’s resistance to CTS increases by orders of magnitude when N122D is combined with Q111R. While not native to the chinchilla lineage, we show that N122D also has the potential to contribute to target-site insensitivity in chinchilla ATP1A1 in the presence of the insM insertion. Surprisingly, N122D does not alter resistance in the absence of the the chinchilla lineage-specific insertion, insM. Our *in silico* modeling analysis shows that insM affects the network of H-bonds between ouabain and several residues within the CTS binding pocket. These bonds are weakened by the removal of insM but can also be strengthened if its removal is combined with N122D.

The wildtype sandgrouse ATP1A1, which has R111, exhibits a level of CTS resistance on par with a CTS-sensitive ATP1A1 paralog (ATP1A1S) of neotropical grass frogs engineered to carry the Q111R mutation (Mohammadi et al. 2022). However, unlike the frog, in the sandgrouse ATP1A1, there is no difference in levels of resistance between amino acid states Q111 and R111. We found instead that sandgrouse ATP1A1 retains resistance with R111 alone or with insD alone; resistance is only lost when both states are reverted. Our protein structure models reveal that mutating R111Q in the absence of insD leads to the formation of multiple strong hydrogen bonds, enabling high affinity complex formation. It is important to note that our docking simulations model the biophysical interactions of ouabain within the NKA’s binding cavity, and that it is possible that these amino acid residues influence the protein’s “gatekeeping” function, preventing ouabain from entering the binding pocket.

Given that the wildtype chinchilla ATP1A1 lacks target-site insensitivity, our results also illustrate how epistasis can represent a source of contingency in the evolution of CTS resistance. For example, in the case of chinchilla ATP1A1, the insM mutation influences which amino acid substitutions could potentially contribute to the evolution of target-site insensitivity. If chinchillas or other hystrigonath rodents with insM were subject to selection for CTS resistance, the presence of insM would potentiate the effect of N122D, an amino acid substitution that confers target-site insensitivity in the NKA of other rodents. Thus, although the insM mutation might have been neutral in the ancestor of hystrigonaths, it influences the selection coefficients associated with future mutations in the same protein. Interestingly, other rodents and frogs that have evolved both Q111R and N122D lack an insertion like insM or insD. We speculate that rodents that share these same amino acid states have other amino acid substitutions that serve a compensatory function, similar to the insertions in chinchilla and sandgrouse ATP1A1. The only plausible candidates in the H1-H2 loop are a serine/threonine substitution a position 112 (A119S/T, shared with hystricognaths) and a proline substitution at residue 119, a site that has previously been implicated in compensatory interactions with substitutions at sites 111 and 122 (Karageorgi et al. 2019; Taverner et al. 2019). Similarly, it has been shown that frogs have 10 or fewer additional substitutions that mitigate negative pleiotropic effects caused by Q111R and N122D in a CTS resistant paralog of ATP1A1 (Mohammadi et al. 2021), only two of which occur in the H1-H2 loop (A112S/T shared with hystricognaths and E116D).

Our functional and modeling data reveal potential mechanisms for mitigating negative pleiotropic effects of resistance-conferring mutations and provide insights into the role of indels in protein evolution. A question that remains open is whether these insertions evolved first and were later compensated for by flanking substitutions or whether the flanking substitutions evolved first, thereby producing a permissive background for the insertion to become fixed. It is thought that amino acid insertion and deletion mutations (indels) may have lower fixation probabilities than amino acid point mutations because the former are more likely to disrupt protein function (de la Chaux et al. 2007; Hu and Ng 2012; Montgomery et al. 2013). Indels have been shown to evolve under strong purifying selection and accumulating evidence indicates that point mutations and indels in proteins are interdependent (Tian et al. 2008; Chen et al. 2009; Tóth-Petróczy and Tawfik 2013). Further, protein regions flanking indels have been shown to evolve faster than the rest of the protein, suggesting that indels and associated substitutions may accumulate in a neutral-compensatory manner (Tian et al. 2008; Tóth-Petróczy and Tawfik 2013). We observe a similar pattern of amino acid substitutions correlated with insertions in the hystricognath and sandgrouse clades (Figure 2), whose H1-H2 loop amino acid sequences are markedly divergent from sister taxa. We speculate that the other derived sites (beyond 111 and 122) in this loop also contribute to epistatic effects on CTS resistance and protein activity, and further experimental tests should help dissect these mechanisms.

## Acknowledgements

We thank P. Kowalski, M. Herbertz, and V. Wagschal for their assistance in the laboratory. This study was funded by grants to PA from the National Institutes of Health (R01–GM115523), to JFS from the National Institutes of Health (R01–HL087216) and the National Science Foundation (OIA–1736249), to SD from the Deutsche Forschungsgemeinschaft (Do 517/10-1), and to SM from the Alexander von Humboldt-Stiftung (Mohammadi 2019) and the National Institutes of Health (F32–HL149172).

## Materials and Methods

### Data sources

Sequence data for ATP1A1 of the common chinchilla and 25 related rodents, and of the yellow-throated sandgrouse and 21 related birds were collected from publicly available sources (Table S1). Eighteen of the bird ATP1A1 sequences were obtained from genome assemblies under BioProject PRJNA545868 (Feng et al. 2020). ATP1B1 sequences for *Chinchilla lanigera* (Genbank: XM005398203) and *Pterocles gutturalis* (Genbank: XM010081314) were also collected from publicly available sources. Previously characterized functional properties of two recombinant chinchilla and two recombinant yellow-throated sandgrouse NKAs were obtained from a previous study (Mohammadi et al. 2022) (see Table S2).

### Phylogenetic tree inference

Nucleotide sequences representing ATP1A1 cDNA for rodents and birds were aligned separately using the MAFFT function in Geneious Prime v 2021.0.3 (Biomatters Ltd). The alignments were analyzed with maximum likelihood (ML) inference using IQ-TREE v 2.1.2 (Minh et al. 2020). The IQ-TREE analysis was run using codon-based sequences (-st) and the best-fit model for each partition with 1000 ultrafast bootstrap (UFB) replicates. All IQ-TREE analyses were performed using the CIPRES Science Gateway online server (Miller et al. 2012). To describe ancestral states and substitutions, we used standardized numbering of residues based on the sheep (*Ovis ares)* ATP1A1 sequence (Genbank: NC019458.2) minus 5 residues from the 5’ end.

### Construction of expression vectors

To supplement previous data (Mohammadi et al. 2022), we created four additional mutagenized versions of ATP1A1 sequences (Invitrogen™ GeneArt) that were codon optimized for *Spodoptera frugiperda* (Table 1). Newly generated plasmid constructs were deposited at Addgene repository under accession numbers 191226-191229. We used recombinant NKA protein constructs for chinchilla and sandgrouse generated by Mohammadi et al. (Mohammadi et al. 2022). First, ATP1B1 genes were inserted into pFastBac ™ Dual Expression Vector (Thermo Scientific™; Cat#10712024) at the p10 promoter with XhoI and PaeI (FastDigest; Thermo Scientific™; Cat#FD0694 and Cat#FD0593, respectively) and confirmed by sequencing. The ATP1A1 genes were inserted at the P_PH_ promoter of vectors already containing the corresponding ATP1B1 genes using In-Fusion^®^ HD Cloning Kit (Takara Bio; Cat#638910) and confirmed by sequencing. All resulting vectors had the ATP1A1 gene under the control of the P_PH_ promoter and a ATP1B1 gene under the p10 promoter. The resulting two vectors were then subjected to site-directed mutagenesis (QuickChange II XL Site-Directed Mutagenesis Kit; Agilent Technologies, La Jolla, CA, USA; Cat#200521) to introduce the amino acid codons of interest.

### Generation of recombinant viruses and transfection into Sf9 cells (Spodoptera frugiperda)

*Escherichia coli* DH10bac cells harboring the baculovirus genome (bacmid) and a transposition helper vector (Thermo Fisher Scientific™; Cat#10361012) were transformed according to the manufacturer’s protocol with expression vectors containing the different gene constructs. Recombinant bacmids were selected through PCR screening, grown, and isolated. Subsequently, Sf9 cells (4 x 10^5^ cells*ml) in 2 ml of Insect-Xpress medium (Lonza; Cat#BE12-730P10) were transfected with recombinant bacmids using Cellfectin II reagent (Gibco-Thermo Fisher Scientific™; Cat#10362100). After a three-day incubation period, recombinant baculoviruses were isolated (P1) and used to infect fresh Sf9 cells (1.2 x 10^6^ cells*ml) in 10 ml of Insect-Xpress medium with 15 mg/ml gentamycin (Roth; Cat#0233.1) at a multiplicity of infection of 0.1. Five days after infection, the amplified viruses were harvested (P2 stock).

### Preparation of Sf9 membranes

For production of recombinant NKA, Sf9 cells were infected with the P2 viral stock at a multiplicity of infection of 1e3. The cells (1.6 x 10^6^ cells*ml) were grown in 50 ml of Insect-Xpress medium with 15 mg/ml gentamycin at 27°C in 500 ml flasks (Dalla et al. 2017). After 3 days, Sf9 cells were harvested by centrifugation at 3,000 x g for 10 min. The cells were stored at −80 °C, and then resuspended at 0 °C in 15 ml of homogenization buffer (0.25 M sucrose, 2 mM EDTA, and 25 mM HEPES/Tris; pH 7.0). The resuspended cells were sonicated at 60 W (Sonopuls 2070; Bandelin Electronic Company, Berlin, Germany) for three 45 s intervals at 0 °C. The cell suspension was then subjected to centrifugation for 30 min at 10,000 x g (J2-21 centrifuge, Beckmann-Coulter, Krefeld, Germany). The supernatant was collected and further centrifuged for 60 min at 100,000 x g at 4 °C (Ultra-Centrifuge L-80, Beckmann-Coulter) to pellet the cell membranes. The pelleted membranes were washed twice and resuspended in ROTIPURAN^®^ p.a., ACS water (Roth; Cat#HN68.2) and stored at −20 °C. Protein concentrations were determined by Bradford assays using bovine serum albumin as a standard. Three biological replicates were produced for each NKA.

### Verification by SDS-PAGE/western blotting

For each biological replicate, 10 ug of protein were solubilized in 4x SDS-polyacrylamide gel electrophoresis sample buffer and separated on SDS gels containing 10% acrylamide. Subsequently, they were blotted on nitrocellulose membrane (Roth; Cat#HP42.1). To block non-specific binding sites after blotting, the membrane was incubated with 5% dried milk in TBS-Tween 20 for 1 h. After blocking, the membranes were incubated overnight at 4 °C with the primary monoclonal antibody α5 (Developmental Studies Hybridoma Bank, University of Iowa, Iowa City, IA, USA; RRID:AB_2166869). Because only membrane proteins were isolated from transfected cells, detection of the α subunit also indicates the presence of the β subunit. The primary antibody was detected using a goat-anti-mouse secondary antibody conjugated with horseradish peroxidase (Dianova, Hamburg, Germany; Cat#115-035-003; RRID:AB_2617176). The staining of the precipitated polypeptide-antibody complexes was performed by addition of 60 mg 4-chloro-1 naphtol (Merck/Sigma-Aldrich; Cat#C8890) in 20 ml ice-cold methanol to 100 ml phosphate buffered saline (PBS) containing 60 ul 30% H2O2. See Figure S1.

### Ouabain inhibition assay

To determine the sensitivity of each NKA construct against the water-soluble cardiotonic steroid, ouabain (Ouabain octahydrate 96%; Acrōs Organic; Cat#AC161730010s), 100 ug of each protein was pipetted into each well in a nine-well row on a 96-well flat-bottom microplate containing stabilizing buffers (see buffer formulas in (Petschenka et al., 2013)). Each well in the nine-well row was exposed to exponentially decreasing concentrations (10^-3^ M, 10^-4^ M, 10^-5^ M, 10^-6^ M, 10^-7^ M, 10^-8^ M, dissolved in distilled H2O) of ouabain, HPLC-grade water only (experimental control), and a combination of an inhibition buffer lacking KCl and 10^-2^ M ouabain (ouabain octahydrate 96%; Acrōs Organic; Cat#AC161730010s) to measure background protein activity (see (Petschenka et al., 2013)). The proteins were incubated at 37°C and 200 rpms for 10 minutes on a microplate shaker (BioShake iQ; Quantifoil Instruments, Jena, Germany; Cat#1808-0506). Next, ATP (Adenosin-5-triphosphat Bis-(Tris)-salt hydrate; Merck/Sigma-Aldrich; CAS#102047-34-7) was added to each well and the proteins were incubated again at 37°C and 200 rpms for 20 minutes. The activity of Na^+^/K^+^-ATPases following ouabain exposure was determined by quantification of inorganic phosphate (Pi) released from enzymatically hydrolyzed ATP. Reaction Pi levels were measured according to the procedure described by (Taussky and Shorr 1953) (see (Petschenka et al. 2013)). All assays were run in duplicate and the average of the two technical replicates was used for subsequent statistical analyses. Absorbance for each well was measured at 650 nm with a plate absorbance reader (BioRad Model 680 spectrophotometer and software package).

### ATP hydrolysis assay

To determine the functional efficiency of different Na^+^/K^+^-ATPase constructs, we calculated the amount of Pi hydrolyzed from ATP per mg of protein per minute. The measurements were obtained from the same assay as described above. In brief, absorbance from the experimental control reactions, in which 100 ug of protein was incubated without any inhibiting factors (i.e., ouabain or buffer excluding KCl), were measured and translated to mM Pi from a standard curve that was run in parallel (1.2 mM Pi, 1 mM Pi, 0.8 mM Pi, 0.6 mM Pi, 0.4 mM Pi, 0.2 mM Pi, 0 mM Pi). Raw assay data available on Dryad at https://doi.org/10.5061/dryad.ngf1vhhxc.

### Statistical analyses

Background phosphate absorbance levels from reactions with inhibiting factors were used to calibrate phosphate absorbance in wells measuring ouabain inhibition and in the control wells (Petschenka et al. 2013). For ouabain sensitivity measurements, calibrated absorbance values were converted to percentage non-inhibited Na^+^,K^+^-ATPases activity based on measurements from the control wells (Petschenka et al. 2013). These data were plotted and log_10_ IC_50_ values were obtained for each biological replicate from nonlinear fitting using a four-parameter logistic curve, with the top asymptote set to 100 and the bottom asymptote set to zero. Curve fitting was performed with the nlsLM function of the minipack.lm library in R. For comparisons of recombinant protein activity, the calculated Pi concentrations of 100 ug of protein assayed in the absence of ouabain were converted to nmol Pi/mg protein/min (Table S2). IC_50_ values were log_10_-transformed prior to analysis to better meet the assumptions of normality and homogeneity of variance. We used a 2-way ANOVA to assess interaction effects of point mutations and insertion and followed with a Tukey’s test to identify significant differences between the different recombinant proteins (Table S3; Levene’s Test for Homogeneity of Variance for chinchilla IC_50_: F_3,8_= 0.3525 p=0.7888 and protein activity: F_3,8_=0.1622 p=0.9188 and for sandgrouse IC_50_: F_3,8_=0.0243 p=0.9945 and protein activity: F_3,8_=0.4561 p=0.7203). All statistical analyses were implemented in R. Data were plotted using the ggplot2 package in R.

### Homology modelling and *in silico* mutagenesis

The structures of the chinchilla (*Chinchilla lanigera)* and sandgrouse (*Pterocles gutturalis*) NKAs are not available in the protein data bank (PDB) and were thus obtained via homology modelling. The template structures required to perform homology modeling were searched using the BLAST search tool (Altschul et al. 1997) implemented in PyMod 3 (Janson and Paiardini 2021). Crystal structure of high affinity NKA from *Sus scrofa* (PDB ID: 7DDJ with 94.46% and 85.44% sequence identities with chinchilla and sandgrouse respectively) was used as template due to its high homology and higher resolution.

Alignment of template and target sequences for homology modeling was performed using MUSCLE (Edgar 2004) software via PyMod 3 graphical interface. Homology models of the structures were performed using Modeller (Sali and Blundell 1993) implemented in PyMod 3. In addition, regions with low DOPE scores (Shen and Sali 2006), including the loops in the binding region, were further refined after initial modeling using Modeller. Following standard procedures, ligand molecules and N-terminal amino acids were deleted and disulfide bonds were patched for more accurate modeling (Gray et al. 2003). The modeled structures were energetically minimized with the 1000-step Steepest Descent algorithm using the AMBER99SB-ILDN force field (Lindorff-Larsen et al. 2010) in the OpenMM toolkit (Eastman et al. 2017). Finally, the required point mutations were performed using PyMol (Schrödinger 2015) mutation wizard, where the romaters with a probability greater than 20% are chosen. After mutations, minimization is performed under the same conditions for another 1000 steps using the OpenMM toolkit to optimize amino acid side chain orientations.

### Molecular docking

Docking calculations for the modeled ATPase structures were performed using Autodock Vina 1.1.2 (Trott and Olson 2010). For docking simulations, ouabain (OBN) ligand molecule was extracted from the NKA-ouabain co-crystallized structure (PDB ID: 7DDJ) from Protein Data Bank (PDB) (Berman et al. 2002). Hydrogen atoms were added and the point charges were corrected using AutoDock Tools (ADT) graphical interface software included in MGLTools 1.5.7 (Sanner 1999). A grid box of dimensions 35×35×40 Å was constructed to include the binding pocket of the ligand for all docking experiments based on the co-crystal structure of ouabain and ATPase complex (PDB ID: 7DDJ) and exhaustiveness value was taken as 10. In addition, interacting residues were selected from the ATPase and OBN complex co-crystal structure (PDB ID: 7DDJ) using LIGPLOT (Wallace et al. 1995) (Fig. 3E-F). In particular, residues involved in hydrogen bonding (Q111, Q119, E312 and T797) were selected to be flexible in the docking process (Table S4) (Ravindranath et al. 2015). For each docking calculation, 10 repetitions were performed and poses with lowest docking scores (low scores correspond to best structures—the ones with highest affinity) were extracted (Fig. 3, Fig. S2). The pose with the lowest docking score corresponds to the best binding ligand. PyMOL was used for visual inspection of the docked structures and Discovery Studio was used for hydrogen binding determinations.

## References

Agrawal AA, Petschenka G, Bingham RA, Weber MG, Rasmann S. 2012. Toxic cardenolides: chemical ecology and coevolution of specialized plant-herbivore interactions. New Phytol. 194:28–45.

Altschul SF, Madden TL, Schäffer AA, Zhang J, Zhang Z, Miller W, Lipman DJ. 1997. Gapped BLAST and PSI-BLAST: a new generation of protein database search programs. Nucleic Acids Res.25:3389–3402.

Aperia A. 2007. New roles for an old enzyme: Na, K-ATPase emerges as an interesting drug target. J. Intern. Med. 261:44–52.

Bagrov AY, Shapiro JI, Fedorova OV. 2009. Endogenous cardiotonic steroids: physiology, pharmacology, and novel therapeutic targets. Pharmacol. Rev. 61:9–38.

Berman HM, Battistuz T, Bhat TN, Bluhm WF, Bourne PE, Burkhardt K, Feng Z, Gilliland GL, Iype L, Jain S. 2002. The protein data bank. Acta Crystallogr. D Biol. Crystallogr. 58:899–907.

Blanco G, Mercer RW. 1998. Isozymes of the Na-K-ATPase: heterogeneity in structure, diversity in function. Am. J. Physiol.-Ren. Physiol. 275:>F633–F650.

Brodie ED. 2009. Toxins and venoms. Curr. Biol. 19:R931–R935.

de la Chaux N, Messer PW, Arndt PF. 2007. DNA indels in coding regions reveal selective constraints on protein evolution in the human lineage. BMC Evol. Biol. 7:1–12.

Chen J-Q, Wu Y, Yang H, Bergelson J, Kreitman M, Tian D. 2009. Variation in the Ratio of Nucleotide Substitution and Indel Rates across Genomes in Mammals and Bacteria. Mol. Biol. Evol.26:1523–1531.

Dalla S, Baum M, Dobler S. 2017. Substitutions in the cardenolide binding site and interaction of subunits affect kinetics besides cardenolide sensitivity of insect Na, K-ATPase. Insect Biochem. Mol. Biol. 89:43–50.

Dobler S, Dalla S, Wagschal V, Agrawal AA. 2012. Community-wide convergent evolution in insect adaptation to toxic cardenolides by substitutions in the Na, K-ATPase. Proc. Natl. Acad. Sci.109:13040–13045.

Dobler S, Wagschal V, Pietsch N, Dahdouli N, Meinzer F, Romey-Glüsing R, Schütte K. 2019. New ways to acquire resistance: imperfect convergence in insect adaptations to a potent plant toxin. Proc. Biol. Sci. 286:20190883.

Eastman P, Swails J, Chodera JD, McGibbon RT, Zhao Y, Beauchamp KA, Wang L-P, Simmonett AC, Harrigan MP, Stern CD. 2017. OpenMM 7: Rapid development of high performance algorithms for molecular dynamics. PLoS Comput. Biol. 13:e1005659.

Edgar RC. 2004. MUSCLE: multiple sequence alignment with high accuracy and high throughput. Nucleic Acids Res. 32:1792–1797.

Feng S, Stiller J, Deng Y, Armstrong J, Fang Q, Reeve AH, Xie D, Chen G, Guo C, Faircloth BC. 2020. Dense sampling of bird diversity increases power of comparative genomics. Nature 587:252–257.

Gong LI, Bloom JD. 2014. Epistatically interacting substitutions are enriched during adaptive protein evolution. PLoS Genet. 10:e1004328.

Gray JJ, Moughon S, Wang C, Schueler-Furman O, Kuhlman B, Rohl CA, Baker D. 2003. Protein–protein docking with simultaneous optimization of rigid-body displacement and side-chain conformations. J. Mol. Biol. 331:281–299.

Groen SC, Whiteman NK. 2021. Convergent evolution of cardiac-glycoside resistance in predators and parasites of milkweed herbivores. Curr. Biol. CB 31:R1465–R1466.

Hu J, Ng PC. 2012. Predicting the effects of frameshifting indels. Genome Biol. 13:1–11.

Hutchinson DA, Mori A, Savitzky AH, Burghardt GM, Wu X, Meinwald J, Schroeder FC. 2007. Dietary sequestration of defensive steroids in nuchal glands of the Asian snake Rhabdophis tigrinus. Proc. Natl. Acad. Sci. U. S. A. 104:2265–2270.

Janson G, Paiardini A. 2021. PyMod 3: a complete suite for structural bioinformatics in PyMOL. Bioinformatics 37:1471–1472.

Kanai R, Cornelius F, Ogawa H, Motoyama K, Vilsen B, Toyoshima C. 2021. Binding of cardiotonic steroids to Na+, K+-ATPase in the E2P state. Proc. Natl. Acad. Sci. 118.

Karageorgi M, Groen SC, Sumbul F, Pelaez JN, Verster KI, Aguilar JM, Hastings AP, Bernstein SL, Matsunaga T, Astourian M, et al. 2019. Genome editing retraces the evolution of toxin resistance in the monarch butterfly. Nature 574:409–412.

Köksoy AA. 2002. Na+, K+-ATPase: a review. J Ank. Med Sch 24:73–82.

Krenn L, Kopp B. 1998. Bufadienolides from animal and plant sources. Phytochemistry 48:1–29.

Kuhl H, Frankl-Vilches C, Bakker A, Mayr G, Nikolaus G, Boerno ST, Klages S, Timmermann B, Gahr M. 2021. An unbiased molecular approach using 3’-UTRs resolves the avian family-level tree of life. Mol. Biol. Evol. 38:108–127.

Laursen M, Gregersen JL, Yatime L, Nissen P, Fedosova NU. 2015. Structures and characterization of digoxin-and bufalin-bound Na+,K+-ATPase compared with the ouabain-bound complex. Proc. Natl. Acad. Sci. U. S. A. 112:1755–1760.

Lindorff-Larsen K, Piana S, Palmo K, Maragakis P, Klepeis JL, Dror RO, Shaw DE. 2010. Improved side-chain torsion potentials for the Amber ff99SB protein force field. Proteins 78:1950–1958.

Marivaux L, Vianey-Liaud M, Jaeger J-J. 2004. High-level phylogeny of early Tertiary rodents: dental evidence. Zool. J. Linn. Soc. 142:105–134.

Marshall BM, Casewell NR, Vences M, Glaw F, Andreone F, Rakotoarison A, Zancolli G, Woog F, Wüster W. 2018. Widespread vulnerability of Malagasy predators to the toxins of an introduced toad. Curr. Biol. CB 28:2194.

McGlothlin JW, Kobiela ME, Feldman CR, Castoe TA, Geffeney SL, Hanifin CT, Toledo G, Vonk FJ, Richardson MK, Brodie Jr ED. 2016. Historical contingency in a multigene family facilitates adaptive evolution of toxin resistance. Curr. Biol. 26:1616–1621.

Miller MA, Pfeiffer W, Schwartz T. 2012. The CIPRES science gateway: enabling high-impact science for phylogenetics researchers with limited resources. In: p. 1–8.

Minh BQ, Schmidt HA, Chernomor O, Schrempf D, Woodhams MD, Von Haeseler A, Lanfear R. 2020. IQ-TREE 2: new models and efficient methods for phylogenetic inference in the genomic era. Mol. Biol. Evol. 37:1530–1534.

Mobasheri A, Avila J, Cózar-Castellano I, Brownleader MD, Trevan M, Francis MJ, Lamb JF, Martín-Vasallo P. 2000. Na+, K+-ATPase isozyme diversity; comparative biochemistry and physiological implications of novel functional interactions. Biosci. Rep. 20:51–91.

Mohammadi S, Gompert Z, Gonzalez J, Takeuchi H, Mori A, Savitzky AH. 2016. Toxin-resistant isoforms of Na+/K+-ATPase in snakes do not closely track dietary specialization on toads. Proc. Biol. Sci. 283:20162111.

Mohammadi S, Herrera-Álvarez S, Yang L, del Pilar Rodríguez-Ordoñez M, Zhang K, Storz JF, Dobler S, Crawford AJ, Andolfatto P. 2022. Constraints on the evolution of toxin-resistant Na,K-ATPases have limited dependence on sequence divergence. PLoS Genet. 18:e1010323.

Mohammadi S, Yang L, Harpak A, Herrera-Álvarez S, Del Pilar Rodríguez-Ordoñez M, Peng J, Zhang K, Storz JF, Dobler S, Crawford AJ, et al. 2021. Concerted evolution reveals co-adapted amino acid substitutions in Na+K+-ATPase of frogs that prey on toxic toads. Curr. Biol. CB 31:2530–2538.e10.

Montgomery SB, Goode DL, Kvikstad E, Albers CA, Zhang ZD, Mu XJ, Ananda G, Howie B, Karczewski KJ, Smith KS, et al. 2013. The origin, evolution, and functional impact of short insertion-deletion variants identified in 179 human genomes. Genome Res. 23:749–761.

Petschenka G, Fandrich S, Sander N, Wagschal V, Boppré M, Dobler S. 2013. Stepwise evolution of resistance to toxic cardenolides via genetic substitutions in the Na+/K+-ATPase of milkweed butterflies (lepidoptera: Danaini). Evol. Int. J. Org. Evol. 67:2753–2761.

Price EM, Lingrel JB. 1988. Structure-function relationships in the Na,K-ATPase alpha subunit: site-directed mutagenesis of glutamine-111 to arginine and asparagine-122 to aspartic acid generates a ouabain-resistant enzyme. Biochemistry 27:8400–8408.

Price EM, Rice DA, Lingrel JB. 1990. Structure-function studies of Na, K-ATPase. Site-directed mutagenesis of the border residues from the H1-H2 extracellular domain of the alpha subunit. J. Biol. Chem. 265:6638–6641.

Ravindranath PA, Forli S, Goodsell DS, Olson AJ, Sanner MF. 2015. AutoDockFR: advances in protein-ligand docking with explicitly specified binding site flexibility. PLoS Comput. Biol. 11:e1004586.

Sali A, Blundell TL. 1993. Comparative protein modelling by satisfaction of spatial restraints. J. Mol. Biol. 234:779–815.

Sallam HM, Seiffert ER, Steiper ME, Simons EL. 2009. Fossil and molecular evidence constrain scenarios for the early evolutionary and biogeographic history of hystricognathous rodents. Proc. Natl. Acad. Sci. U. S. A. 106:16722–16727.

Sanner MF. 1999. Python: a programming language for software integration and development. J. Mol. Graph. Model. 17:57–61.

Schrödinger L. 2015. The PyMOL molecular graphics system, version 1.8.

Shen M-Y, Sali A. 2006. Statistical potential for assessment and prediction of protein structures. Protein Sci. Publ. Protein Soc. 15:2507–2524.

Storz JF. 2016. Causes of molecular convergence and parallelism in protein evolution. Nat. Rev. Genet.17:239–250.

Storz JF. 2018. Compensatory mutations and epistasis for protein function. Curr. Opin. Struct. Biol.50:18–25.

Taussky HH, Shorr E. 1953. A microcolorimetric method for the determination of inorganic phosphorus. J. Biol. Chem. 202:675–685.

Taverner AM, Yang L, Barile ZJ, Lin B, Peng J, Pinharanda AP, Rao AS, Roland BP, Talsma AD, Wei D, et al. 2019. Adaptive substitutions underlying cardiac glycoside insensitivity in insects exhibit epistasis in vivo. eLife 8:e48224.

Tian D, Wang Q, Zhang P, Araki H, Yang S, Kreitman M, Nagylaki T, Hudson R, Bergelson J, Chen J-Q. 2008. Single-nucleotide mutation rate increases close to insertions/deletions in eukaryotes. Nature 455:105–108.

Tóth-Petróczy Á, Tawfik DS. 2013. Protein Insertions and Deletions Enabled by Neutral Roaming in Sequence Space. Mol. Biol. Evol. 30:761–771.

Trott O, Olson AJ. 2010. AutoDock Vina: improving the speed and accuracy of docking with a new scoring function,efficient optimization, and multithreading. J. Comput. Chem. 31:455–461.

Ujvari B, Casewell NR, Sunagar K, Arbuckle K, Wüster W, Lo N, O’Meally D, Beckmann C, King GF, Deplazes E, et al. 2015. Widespread convergence in toxin resistance by predictable molecular evolution. Proc. Natl. Acad. Sci. U. S. A. 112:11911–11916.

Ujvari B, Mun H, Conigrave AD, Bray A, Osterkamp J, Halling P, Madsen T. 2013. Isolation breeds naivety: island living robs Australian varanid lizards of toad-toxin immunity via four-base-pair mutation. Evol. Int. J. Org. Evol. 67:289–294.

Wallace AC, Laskowski RA, Thornton JM. 1995. LIGPLOT: a program to generate schematic diagrams of protein-ligand interactions. Protein Eng. Des. Sel. 8:127–134.

Weinreich DM, Delaney NF, DePristo MA, Hartl DL. 2006. Darwinian evolution can follow only very few mutational paths to fitter proteins. science 312:111–114.

Yang L, Ravikanthachari N, Mariño-Pérez R, Deshmukh R, Wu M, Rosenstein A, Kunte K, Song H, Andolfatto P. 2019. Predictability in the evolution of Orthopteran cardenolide insensitivity. Philos. Trans. R. Soc. Lond. B. Biol. Sci. 374:20180246.

Zhen Y, Aardema ML, Medina EM, Schumer M, Andolfatto P. 2012. Parallel molecular evolution in an herbivore community. Science 337:1634–1637.

