## Supplemental Information for "Epistatic effects between amino acid insertions and substitutions mediate toxin-resistance of vertebrate Na^+^,K^+^-ATPases"

**Table S1.** Sources of sequences used in phylogenetic tree inference.

| Species | Group | Data type and format | Accession |
|---|---|---|---|
| *Ardeotis kori* ATP1A1 | Bird | PRJNA545868 | B10K-CU-031-01 scaffold 1424 |
| *Caloenas nicobarica* ATP1A1 | Bird | PRJNA545868 | OUT-007 scaffold 962 |
| *Centropus bengalensis* ATP1A1 | Bird | PRJNA545868 | B10K-DU-017-21 scaffold 854 |
| *Centropus unirufus* ATP1A1 | Bird | PRJNA545868 | B10K-DU-017-25 scaffold 173 |
| *Ceuthmochares aereus* ATP1A1 | Bird | PRJNA545868 | B10K-CU-031-02 scaffold 3096 |
| *Chlamydotis macqueenii* ATP1A1 | Bird | NCBI reference sequence | XM010121665 |
| *Columba livia* ATP1A1 | Bird | PRJNA545868 | APP-015 scaffold 101 |
| *Columba picui* ATP1A1 | Bird | PRJNA545868 | B10K-DU-021-26 scaffold 962 |
| *Corythaeola cristata* ATP1A1 | Bird | PRJNA545868 | B10K-CU-031-40 scaffold 1516 |
| *Corythaixoides concolor* ATP1A1 | Bird | PRJNA545868 | B10K-DU-011-20 scaffold 545 |
| *Crotophaga sulcirostris* ATP1A1 | Bird | PRJNA545868 | B10K-DU-003-44 scaffold 184 |
| *Cuculus canorus* ATP1A1 | Bird | NCBI reference sequence | XM009556993 |
| *Geococcyx californianus* ATP1A1 | Bird | PRJNA545868 | B10K-CU-031-07 scaffold 43858 |
| *Leptosomus discolor* ATP1A1 | Bird | PRJNA545868 | APP-026 scaffold 4774 |
| *Lophotis ruficrista* ATP1A1 | Bird | PRJNA545868 | B10K-CU-031-23 scaffold 55 |
| *Mesitornis unicolor* ATP1A1 | Bird | NCBI reference sequence | XM010192439 |
| *Patagioenas fasciata* ATP1A1 | Bird | PRJNA545868 | LSYS01009367 |
| *Piaya cayana* ATP1A1 | Bird | PRJNA545868 | B10K-DU-008-47 scaffold 11547 |
| *Pterocles burchelli* ATP1A1 | Bird | PRJNA545868 | B10K-DU-027-49 scaffold 1068 |
| *Pterocles gutturalis* ATP1A1 | Bird | NCBI reference sequence | XM010078905 |
| *Syrrhaptes paradoxus* ATP1A1 | Bird | PRJNA545868 | B10K-DU-003-42 scaffold 12443 |
| *Tauraco erythrolophus* ATP1A1 | Bird | NCBI reference sequence | XM009992334 |
| *Arvicanthis niloticus* ATP1A1 | Rodent | NCBI reference sequence | XM034501132 |
| *Castor canadensis* ATP1A1 | Rodent | NCBI reference sequence | GFFW01000188 |
| *Cavia porcellus* ATP1A1 | Rodent | NCBI reference sequence | EF489488 |
| *Chinchilla lanigera* ATP1A1 | Rodent | NCBI reference sequence | XM005389040 |
| *Cricetulus griseus* ATP1A1 | Rodent | NCBI reference sequence | XM007641365 |
| *Dipodomys ordii* ATP1A1 | Rodent | NCBI reference sequence | XM013024679 |
| *Fukomys damarensis* ATP1A1 | Rodent | NCBI reference sequence | XM01631202 |
| *Grammomys surdaster* ATP1A1 | Rodent | NCBI reference sequence | XM028752972 |
| *Heterocephalus glaber* ATP1A1 | Rodent | NCBI reference sequence | XM004853808 |
| *Ictidomys tridecemlineatus* ATP1A1 | Rodent | NCBI reference sequence | XM005334918 |
| *Jaculus jaculus* ATP1A1 | Rodent | NCBI reference sequence | XM004667522 |
| *Marmota flaviventris* ATP1A1 | Rodent | NCBI reference sequence | XM027940230 |
| *Mastomys coucha* ATP1A1 | Rodent | NCBI reference sequence | XM031374991 |
| *Mesocricetus auratus* ATP1A1 | Rodent | NCBI reference sequence | XM005076521 |
| *Microtus ochrogaster* ATP1A1 | Rodent | NCBI reference sequence | XM005357106 |
| *Mus caroli* ATP1A1 | Rodent | NCBI reference sequence | XM021157466 |
| *Mus musculus* ATP1A1 | Rodent | NCBI reference sequence | BC021496 |
| *Mus Pahari* ATP1A1 | Rodent | NCBI reference sequence | XM021195628 |
| *Nannospalax galili* ATP1A1 | Rodent | NCBI reference sequence | XM008834025 |
| *Octodon degus* ATP1A1 | Rodent | NCBI reference sequence | XM023701723 |
| *Peromyscus leucopus* ATP1A1 | Rodent | NCBI reference sequence | XM028877301 |
| *Peromyscus maniculatus* ATP1A1 | Rodent | NCBI reference sequence | XM006989093 |
| *Rattus norvegicus* ATP1A1 | Rodent | NCBI reference sequence | BC061968 |
| *Rattus rattus* ATP1A1 | Rodent | NCBI reference sequence | XM032897619 |
| *Urocitellus parryii* ATP1A1 | Rodent | NCBI reference sequence | XM026394106 |

**Table S2.** Summary of the ouabain sensitivity and catalytic properties of $Na^{+}$,$K^{+}$-ATPase for each recombinant protein construct. The values represent the mean and standard deviation (SD) of ouabain sensitivity (log $IC_{50}$) and protein activity of three biological replicates. An asterisk indicates that data for that construct was obtained from Mohammadi et al. (Mohammadi et al. 2022).

| Construct Name | ouabain sensitivity (mol/L) mean(log $IC_{50}$) ± SD | protein activity nmol Pi/(mg protein*min) ± SD |
|---|---|---|
| *Chinchilla | -5.198 ± 0.207 | 6.464 ± 0.871 |
| *Chinchilla+N122D | -2.333 ± 0.495 | 3.577 ± 0.526 |
| Chinchilla+N122D–insM | -5.811 ± 0.325 | 6.006 ± 1.229 |
| Chinchilla–insM | -3.908 ± 0.276 | 2.481 ± 1.010 |
| *Sandgrouse | -4.679 ± 0.005 | 15.689 ± 2.187 |
| *Sandgrouse+R111Q | -4.698 ± 0.147 | 17.372 ± 3.568 |
| Sandgrouse+R111Q–insD | -5.461 ± 0.177 | 26.588 ± 1.991 |
| Sandgrouse–insD | -4.428 ± 0.222 | 7.492 ± 0.418 |

**Table S3.** ANOVA table and multiple comparison results from Tukey HSD analysis with Bonferroni-corrected p-values. Data associated with Chinchilla were analyzed separately from Sandgrouse. Significant differences are bolded.

| Chinchilla – $IC_{50}$ | Df | Sum Sq | Mean Sq | F value | P value |
|---|---|---|---|---|---|
| N122D | 1 | 0.695 | 0.695 | 5.928 | **0.04090** |
| InsM | 1 | 3.593 | 3.593 | 30.640 | **5.50e-04** |
| N122D:InsM | 1 | 17.054 | 17.054 | 145.433 | **2.06e-06** |
| Residuals | 8 | 0.938 | 0.117 | | |
| Chinchilla – Protein activity | | | | | |
| N122D | 1 | 0.306 | 0.306 | 0.343 | 0.57424 |
| InsM | 1 | 1.812 | 1.812 | 2.032 | 0.19188 |
| N122D:InsM | 1 | 30.827 | 30.827 | 34.572 | **3.7 e-04** |
| Residuals | 8 | 0.892 | 0.892 | | |
| Sandgrouse – $IC_{50}$ | | | | | |
| R111Q | 1 | 1.2740 | 1.2740 | 33.67 | **4.04e-04** |
| InsD | 1 | 0.4373 | 0.4373 | 11.56 | **9.37e-03** |
| R111Q:InsD | 1 | 0.4373 | 0.4373 | 11.56 | **9.37e-03** |
| Residuals | 8 | 0.3027 | 0.0378 | | |
| Sandgrouse – Protein activity | | | | | |
| R111Q | 1 | 323.8 | 323.8 | 59.822 | **5.56e-05** |
| InsD | 1 | 0.8 | 0.8 | 0.144 | 0.714268 |
| R111Q:InsD | 1 | 227.4 | 227.4 | 42.012 | **1.92e-04** |
| Residuals | 8 | 43.3 | 5.4 | | |

| Tukey HSD comparisons Group 1 | Group 2 | $IC_{50}$ adjusted p-value | Protein activity adjusted p-value |
|---|---|---|---|
| Chinchilla | Chinchilla+N122D | **3.39e-05** | **0.023571** |
| Chinchilla | Chinchilla+N122D–insM | 0.204834 | 0.931170 |
| Chinchilla | Chinchilla–insM | **7.48e-03** | **3.77e-03** |
| Chinchilla+N122D | Chinchilla+N122D–insM | **7.7e-06** | **0.053843** |
| Chinchilla+N122D | Chinchilla–insM | **2.19e-03** | 0.521060 |
| Chinchilla+N122D–insM | Chinchilla–insM | **6.24e-04** | **7.89e-03** |
| Sandgrouse | Sandgrouse+R111Q | 0.383429 | 0.812368 |
| Sandgrouse | Sangrouse+R111Q–insD | **8.47e-04** | **1.95e-03** |
| Sandgrouse | Sandgrouse–insD | 1.000000 | **0.010979** |
| Sandgrouse+R111Q | Sangrouse+R111Q–insD | **5.86e-03** | **5.55e-03** |
| Sandgrouse+R111Q | Sandgrouse–insD | 0.383430 | **3.63e-03** |
| Sangrouse+R111Q–insD | Sandgrouse–insD | **8.47e-04** | **3.81e-05** |

**Table S4**. RMSD of ouabain obtained from docking simulations  from reference systems

| **Homology Models** | **RMSD from co-crystal position in 7DDJ* (Å)** | **RMSD from the best WT docking position (Å)** |
|---|---|---|
| Chinchilla | 1.7 | 0.00 |
| Chinchilla + N122D | 1.54 | 0.34 |
| Chinchilla + N122D - insM | 0.52 | 1.73 |
| Chinchilla- insM | 1.96 | 1.43 |
| Sandgrouse | 1.05 | 0.00 |
| Sandgrouse + R111Q | 1.79 | 1.70 |
| Sandgrouse + R111Q - insD | 0.62 | 0.87 |
| Sandgrouse - insD | 1.58 | 1.83 |

*The resolution of the crystal structure (PDB id: 7DDJ) is 2.90 Å

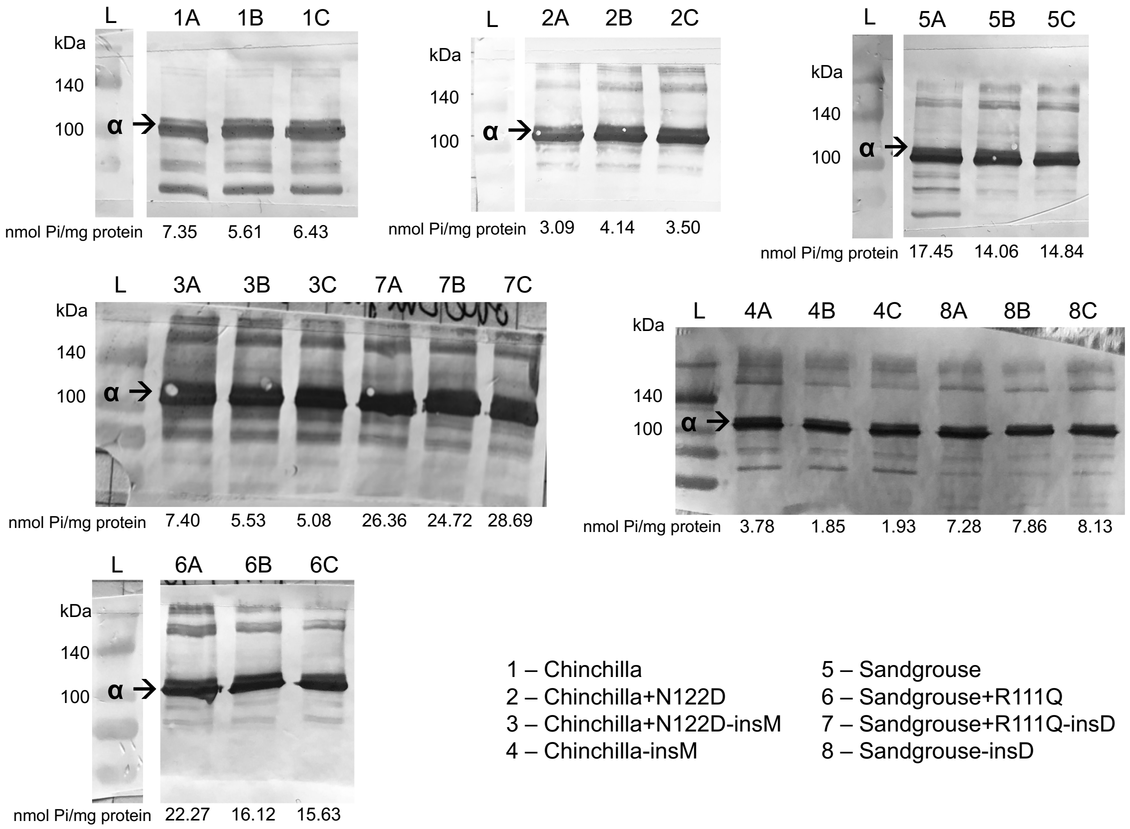


**Figure S1.** Western blot analysis of $Na^+$, $K^+$-ATPase with engineered ATP1A1 (α-subunits) produced in this study. The 110 kDa α-subunits are stained with the α5 monoclonal antibody followed by a horseradish peroxidase conjugated goat antimouse antibody. Samples represents three biological replicates of eight different recombinant $Na^+$, $K^+$-ATPase (Table 1) produced through cell culture. Each panel represents one western blot. The lanes are labeled according to the sample they contained, and the letter "L" indicates the protein ladder. For each western blot, 10 ug of total protein was used. Protein activity levels (nmol*Pi/mg protein) of each biological replicate is indicated under its respective lane. Western blots from constructs 1-2 and 5-6 were obtained from Mohammadi et al. (Mohammadi et al. 2022) – only the ladder and pertinent samples are shown from those western blots.

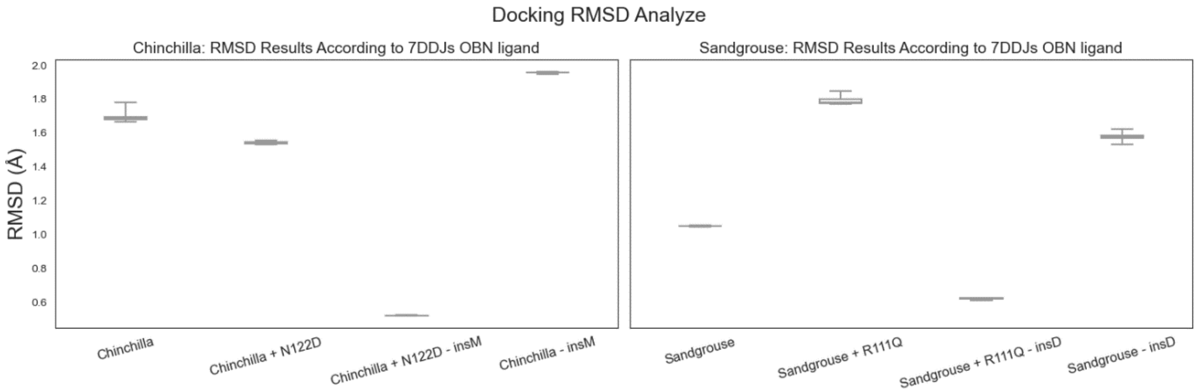


**Figure S2.** The RMSD (root mean square deviation) values of the docked ouabain positions for chinchilla and sandgrouse with respect to the coordinates of ouabain in the co-crystal structure (PDB id: 7DDJ). Each docking simulations are performed with 10 repetitions. The results are shown in standard boxplots, the whiskers represent min/max values and the boxes represents 25$^{th}$ and 75$^{th}$ percentiles. RMSDs were calculated after the transmembrane region of all homology models were aligned to the 7DDJ transmembrane helix.

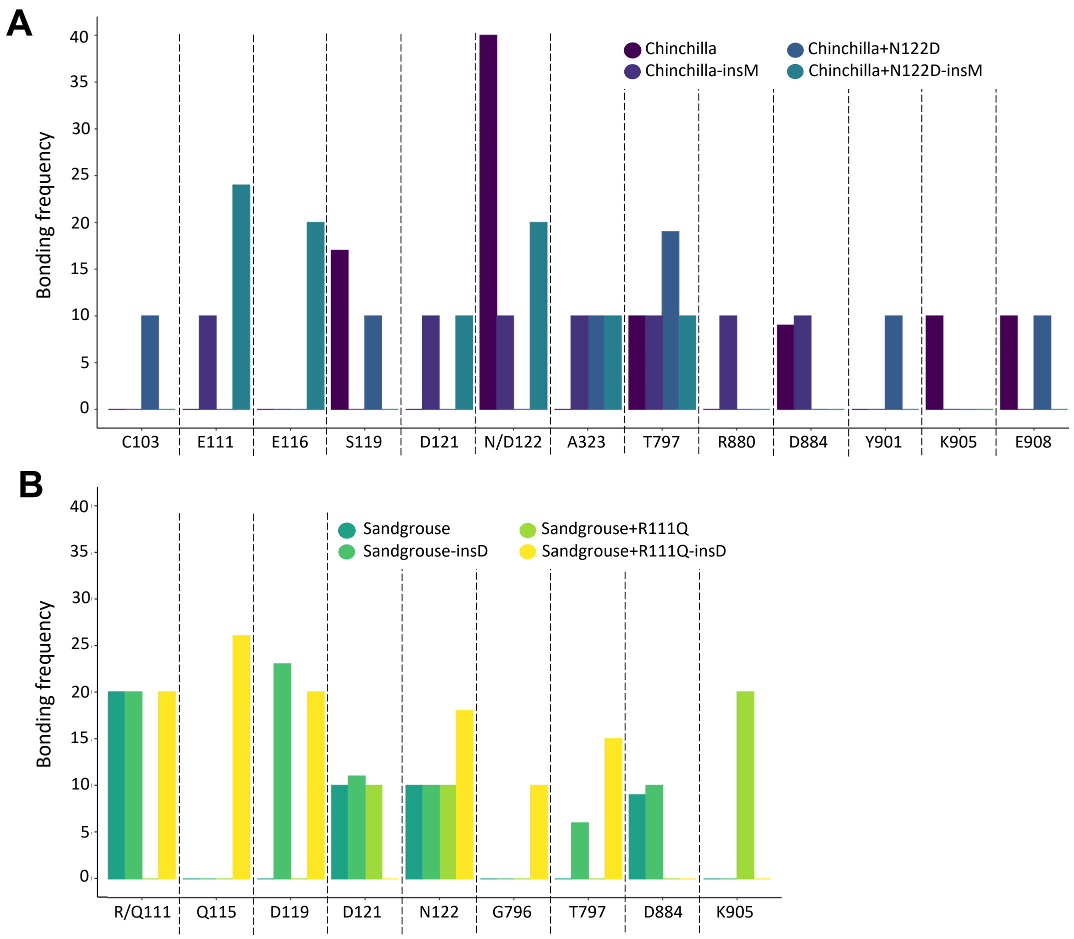


**Figure S3.** The number of instances that hydrogen bonds formed in docking simulations for the four A) chinchilla and B) sandgrouse ATP1A1 structures reveal the likeliness of bonds forming at the given residues. Ten repeats were conducted for each structure. The number of times bonds were formed from ten repeated measurements is plotted on the y-axis. Recombinant protein legend above each graph corresponds to four recombinant protein structures, ordered a-d to identify corresponding bar more easily.

### References


Mohammadi S, Herrera-Álvarez S, Yang L, del Pilar Rodríguez-Ordoñez M, Zhang K, Storz JF, Dobler S, Crawford AJ, Andolfatto P. 2022. Constraints on the evolution of toxin-resistant Na,K-ATPases have limited dependence on sequence divergence. *PLoS genetics* 18:e1010323.
